# ORGANOID AND TISSUE PATTERNING THROUGH PHASE SEPARATION: USE OF A VERTEX MODEL TO RELATE DYNAMICS OF PATTERNING TO UNDERLYING BIOPHYSICAL PARAMETERS

**DOI:** 10.1101/136366

**Authors:** William Waites, Matteo Cavaliere, Élise Cachat, Vincent Danos, Jamie A. Davies

## Abstract

Exactly a century ago, D’Arcy Thompson set an agenda for understanding tissue development in terms of underlying biophysical, mathematically-tractable mechanisms. One such mechanism, discovered by Steinberg in the 1960s, is adhesion-mediated sorting of cell mixtures into homotypic groups. Interest in this phase separation mechanism has recently surged, partly because of its use to create synthetic biological patterning mechanisms and partly because it has been found to drive events critical to the formation of organoids from stem cells, making the process relevant to biotechnology as well as to basic development. Here, we construct quantitative model of patterning by phase separation, informed by laboratory data, and use it to explore the relationship between degree of adhesive difference and speed, type and extent of resultant patterning. Our results can be used three ways; to predict the outcome of mixing cells with known properties, to estimate the properties required to make some designed organoid system, or to estimate underlying cellular properties from observed behaviour.

## 1. Introduction

D’Arcy Thompson sought to explain patterns and forms in living organisms as emergent properties of underlying physical events (Thompson, 1917, 1945). Inspired by pioneering synthetic biologists such as Stephane Leduc (Leduc, 1912), Thompson drew analogies between developing embryos and physicochemical systems such as convection cells and diffusion patterns and proposed that they may use similar underlying mechanisms. A century of embryological progress Entwicklungsmechanik, genetics, cell and molecular biology (see Hamburger, 1988; Keller, 2002, for reviews) has shown the mechanisms of development to be more intricate and complicated than simple convection, diffusion and viscous mixing. Nevertheless, the point on which D’Arcy Thompson insisted in the introduction to *On Growth and Form*, that anatomical complexity emerges from the principles of physics and chemistry with no need for vitalistc mysticism, remains at the heart of experimental embryology.

An aspect of histogenic behaviour that has long been explained in thermodymanic terms, that would have been understood well by Thompson, is the sorting of cells with different adhesive properties. Steinberg established that, in a cell mixture in which cell movement is unconstrained, cells with stronger mutual adhesion will accumulate in the core (Steinberg, 1963). The importance of adhesion itself to this phase separation was later demonstrated by mixing two versions of the same cell line, engineered to express different amounts of the same adhesion molecule (Steinberg and Takeichi, 1994). Steinberg’s thermodynamic explanation was that the system minimizes free energy by minimizing the number of unbound (‘wasted’) adhesion molecules (Steinberg, 1962). Some aspects of ‘cell biology’, such as actin-myosin contraction, are needed for adhesion-mediated cell sorting to work (reviewed by Davies, 2013, pp. 171-176) but sorting behaviour is still predictable on thermodynamic grounds alone (Glazier and Graner, 1993).

Two pieces of recent work demonstrate a role for phase separation not only in tissue development but also in regenerative technologies. One is the spontaneous organization of mixed stem cells to produce organoids (e.g. kidney): in such organoids epithelial stem cells sort out to make epithelial tubules and mesenchymal stem cells organize around them (Unbekandt and Davies, 2010): this self-organization uses adhesion-mediated phase separation (Lefevre et al., 2017). The other is the construction of a synthetic biological system that uses constrained, adhesion-mediated phase separation to produce animal coat-like patterns in cell monolayers (Cachat et al., 2016).

Modelling the parameters of self-organization using adhesion-mediated phase separation is therefore potentially useful for organoid production and tissue synthesis as well as basic developmental biology. In this paper we show that there is a class of physically realistic models, namely *vertex models*, that can be used to generate patterns similar to those produced *in vitro* in Cachat et al. (2016). We demonstrate two techniques for quantifying the degree to which a pattern is present, both applicable to wet-lab studies and to simulations. Having established that vertex models match wet-lab data well, we explore the landscape of adhesion and show that the degree of patterning can be controlled by adjusting the adhesive strength on homotypic and heterotypic cell contacts.

## 2. Results and Discussion

### 2.1. Modelling strategy

We have chosen to use a *vertex model*, which represents each cell in a confluent monolayer as having a certain area and several edges where it meets other cells. This framing is attractive because the objects of interest, cells, are first-class entities and are described in terms of biologically relevant basic physical and geometrical properties (Farhadifar et al., 2007). This contrasts with models that define cells as a collection of contiguous positions on an arbitrary lattice (Glazier and Graner, 1993).

Edges of all cells together form a network (‘graph’) that describes the *topology* of the tissue. The topological properties of this kind of structure are not unique to biological cells and arise *intra alia* with soap bubbles, metal grains and cracks in ceramic glazing (Smith, 1952). With some fairly mild assumptions, namely that the cells are tightly packed, and that exactly three edges come together at each vertex, there are exactly two changes,

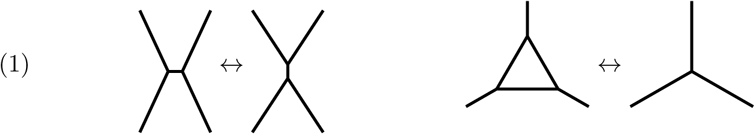

that can generate all possible planar topologies (Weaire and Rivier, 1984). The first is known as the neighbour swap transition or *T1*, and the second is cell extrusion or *T2*. If the latter is not reversible, a third transition is necessary to capture mitosis for modelling proliferation, which we are not concerned with here. The forces governing these transformations were discussed in *On Growth and Form* (Thompson, 1945, pp465-481).

We model the mechanical aspects of the cell monolayer using the potential described in Farhadifar et al. (2007) (see Methods, Section 3.2 and Supplementary Text, Section A.1), an energy cost arising from the competing effects of internal pressure and surface tension. The cost of a given length of edge depends on whether it is in contact with an identical cell or one with different adhesive properties; the cost of an edge *i* is represented by the parameter Λ_*i*_, the value of which is different in homotypic and heterotypic contacts.

As our goal is to find equilibrium solutions for the configuration of the tissue from some arbitrary starting point, integrating the equations of motion directly would lead to oscillation as excess potential energy is converted to kinetic energy that carries the system past equilibrium. Following Mirams et al. (2013) we use a velocity-dependent dissipative force — friction or drag — to remove excess energy from the system (see Supplementary Text, Section A.2). We also constantly perturb the system by a random small amount of ‘noise’ similar to Brownian motion Osborne et al. (2017) (see Methods, Section 3.3).

We use two measures for pattern development, (i) spatial frequency spectral distance and (ii) neighbour entropy; their derivation is explained in Methods. All of our models, real and simulated, begin with two cell types differing only in their adhesive properties.

### 2.2. Development of patterns by phase separation in our model and in real cells

In the real phase separation-driven patterning system of Cachat et al. (2016), cells identical except for carrying either the E-cadherin (‘E-cells’) or Pcadherin (‘P-cells’) version of the construct, mixed randomly to form a confluent monolayer before the construct was activated. Then a small molecule (tetracycline or Tet) was used to induce expression of the cadherin. In the absence of induction, or if all cells were of the same type, no patterning took place. When a 50/50 mix of E-cells and P-cells was induced, they underwent phase separation to form a pattern Fig 1A-D (images obtained from the dataset obtained for Cachat et al. (2016)). If we begin our model with a random 50/50 mix of two cell types, edges of which have different costs according to whether they are homotypic or heterotypic, it generates patterns that are, at least by eye, very similar to the ones produced by the real cells (Fig 1E-F).

**Figure 1.**
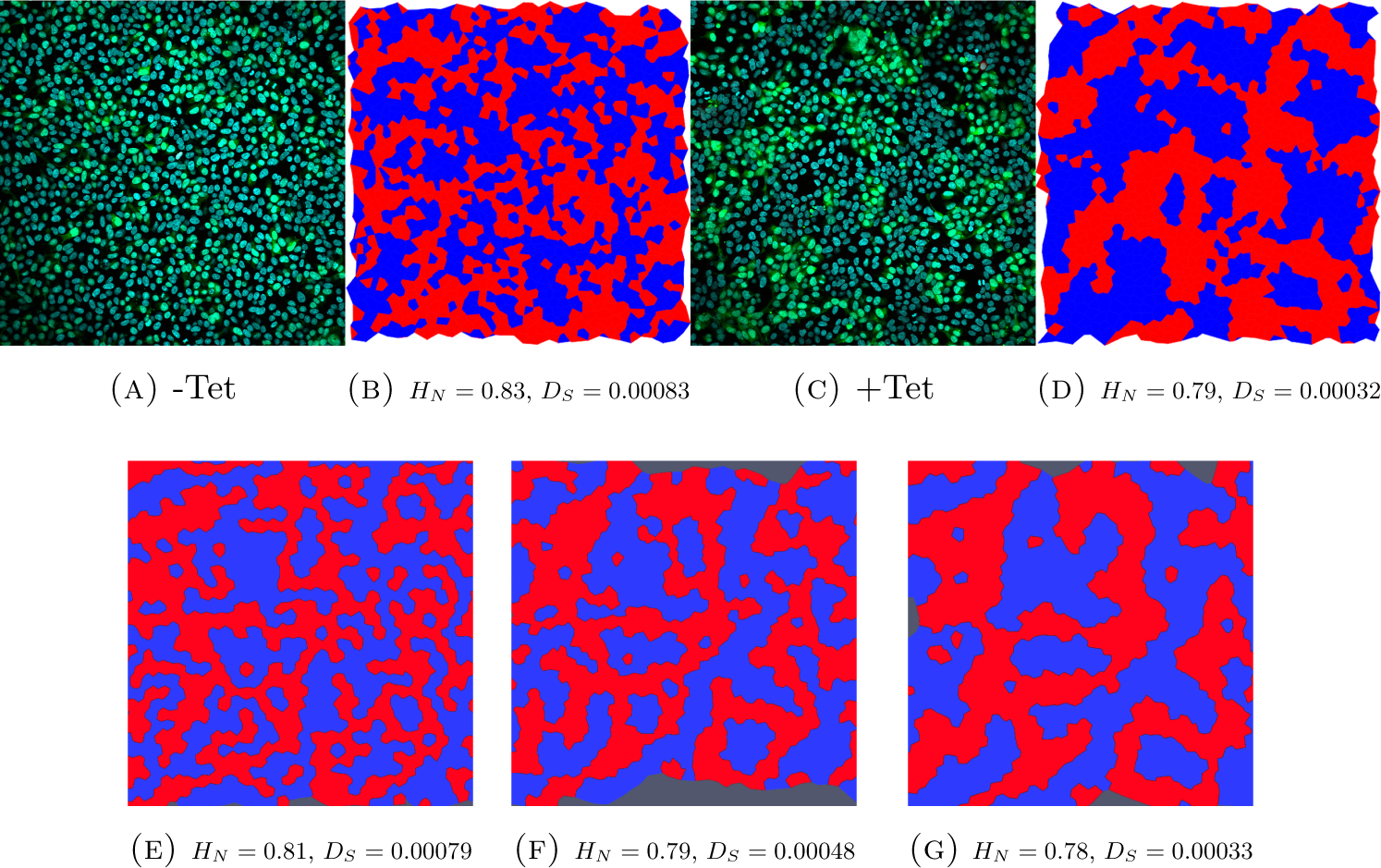
Top row: experimental data. Raw confocal images of 50/50 mixtures of E-cells and P-cells are shown in (A) and (C): all nuclei are stained with DAPI (blue) while only nuclei of E-cells appear green. (A) shows mixtures without induction of adhesion molecule expression and (C) shows mixtures 24h after induction, with patterning of green cells clearly visible. (B) and (D) show segmented images of (A) and (C), E-cells are navy blue in the render used for analysis. Bottom row: simulated data. (E) shows a near-initial state, while (F) and (G) show the system after it has produced a pattern. The character of the pattern in the simulation (F) and (G) is similar to that of the real cells (D), and the neighbour entropy and spectral distance (*H*_*N*_, and *D*_*S*_ shown beneath the images) are respectively identical. Confocal images are 634 *µ*m across and from data published in (Cachat et al., 2016). Simulation data cropped to match the size and resolution of experimental data (see Section 3.5).

To measure the degree of patterning shown, we segmented the images (real or computer-generated: see Supplementary Materials, Sections A.3 and A.4 for more detail) and calculated two measures of patterning.

The basis for our spatial frequency spectral distance measure (*D*_*S*_) is shown in Fig 2: in both the real cells and the computer model, the spectrum of the unpatterned cells displays a greater energy density at a variety of frequencies, with “shoulders” around a central peak (Fig 2C). After patterning the spectrum is much more sharply peaked, with the shoulders nearly gone, and the spectral distance to patterned reference image, *D*_*S*_ falling from 0.0008 to 0.0003.

**Figure 2.**
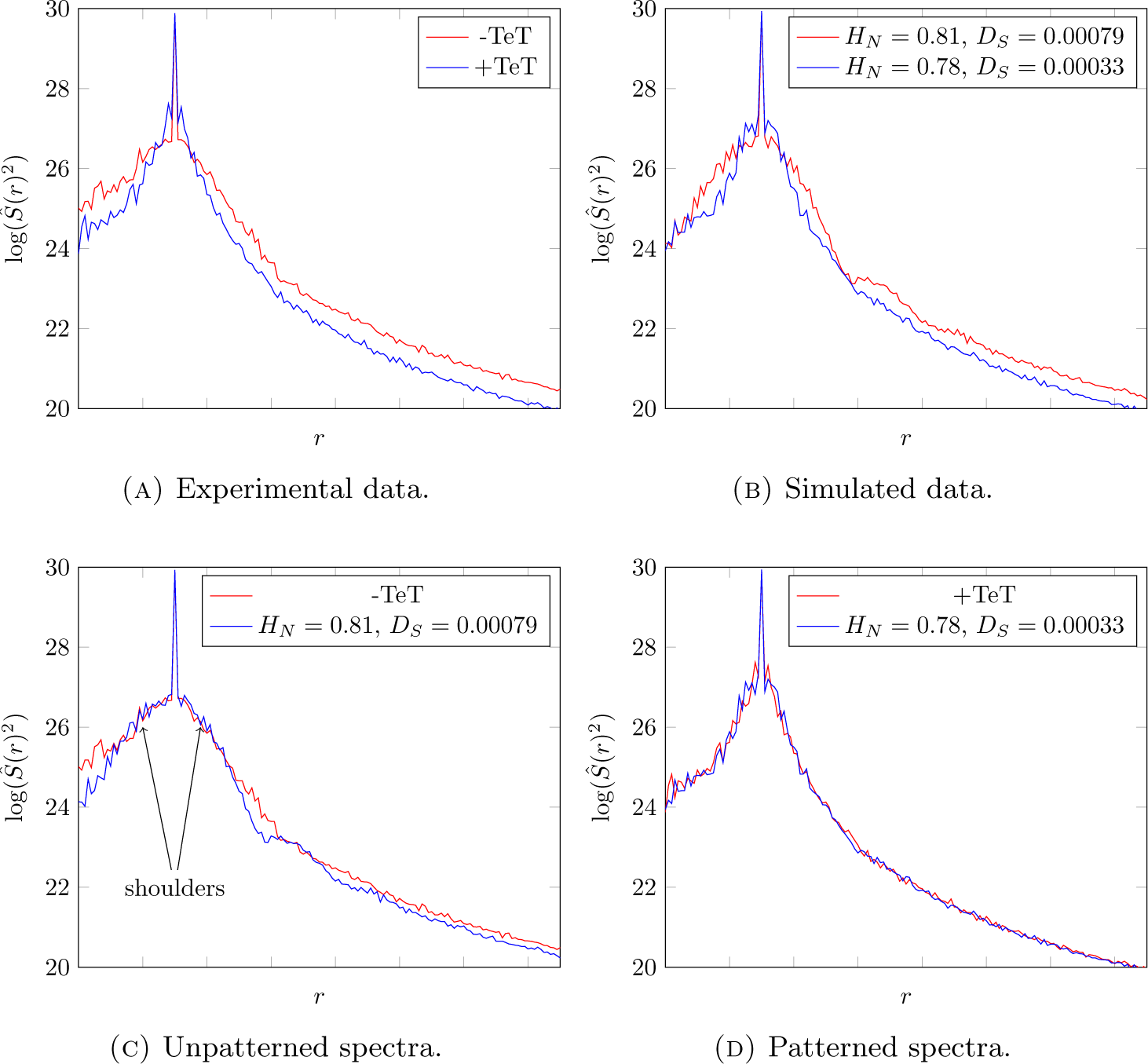
One dimensional power spectra of tissue images. These spectra correspond directly to those of Fig 1. The top row compares spectra for patterned and unpatterned images within each data set, experimental and simulated. The bottom row compares patterned and unpatterned images across data sets. The spectral distance measure is the distance between these (normalised) curves, and (D) in particular suggests that this measure captures the pattern well.

Our other measure, neighbour entropy (*H*_*N*_), also falls from 0.81 to 0.78 (*H*_*N*_ is a sensitive measure and a change of 0.02 represents the difference between pattern and no pattern: see also Fig 3).

**Figure 3.**
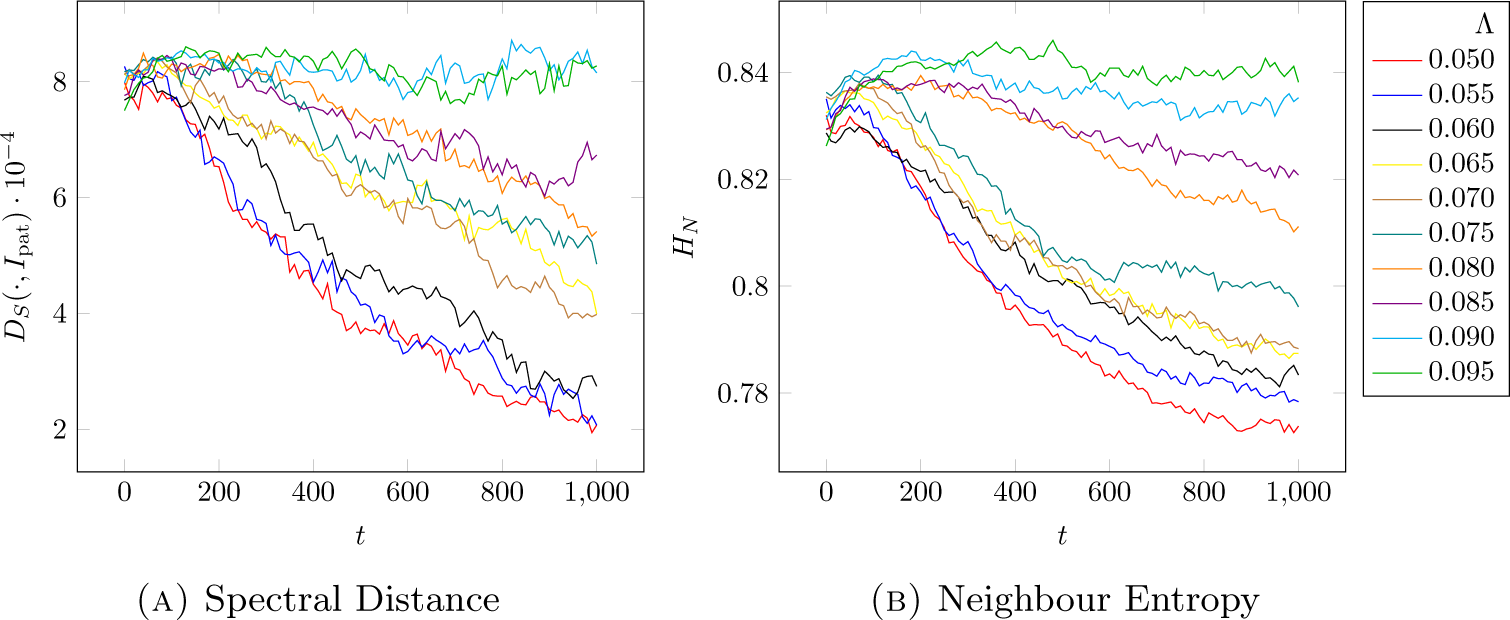
Spectral distance and neighbour entropy for traces for various values of Λ_*PP*_ = Λ_*EE*_. In all cases the heterotypic value is Λ_*P E*_ = Λ_*EP*_ = 0.096.

### 2.3. Analysis of the influence of relative adhesion strengths on patterning

Having shown that the model generates patterns visually similar to those exhibited by real cells, and that this similarity is quantifiable, we have used the model to explore the influence of biophysical parameters on pattern development.

The key parameter is edge cost, Λ_*i*_. Running the simulations from a random mix of two cell types, with the same edge cost for heterotypic contacts, Λ_*PE*_, but different values for homotypic, Λ_*EE*_ = Λ_*PP*_, developed patterns at different rates (Fig 3A). Where the heterotypic edge cost was similar to homotypic (green and blue lines in Fig 3A), patterning was not seen. The lower the homotypic cost, the faster the pattern formed. Similar trends were seen with both measures (Fig 3A,B) although there were some detailed differences in how the lines clustered, especially for intermediate values of homotypic cost.

If the multiple lines shown in Fig 3A,B are each approximated to a straight line by linear regression, the gradient of each can be calculated to allow the plotting of rate of pattern formation against the homotypic edge cost. The results (Fig 4) show that the speed with which a pattern develops, with heterotpyic edge cost held fixed, depends on the cost of the homotypic edge; the greater the difference between homotypic and heterotypic costs, the quicker patterning happens.

**Figure 4.**
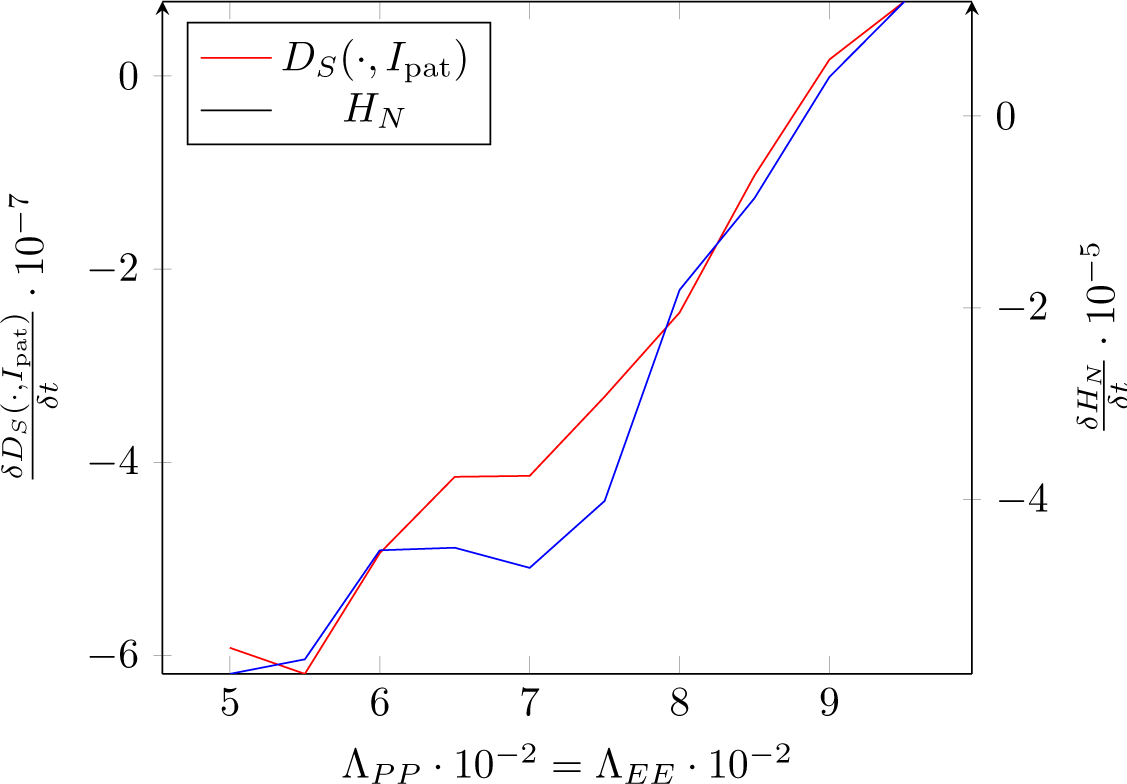
Average rate of change of pattern measures compared to homotypic edge cost.

We also ran a series of simulations holding the homotypic edge cost fixed at the value for quickest patterning (Λ_*P E*_ = Λ_*EP*_ = 0.05) and varying the heterotypic cost: patterning again happened most rapidly when their difference was greatest (see Supplementary Materials, Section B.1). In all cases, because these simulations were not done with periodic boundary conditions, we had the cost of a cell edgeat the edge of the tissue the same as a homotypic edge, that is, Λ_*x·*_ = Λ_*xx*_ for *x ∊ P, E*.

### 2.4. Causal Explanation of Patterning through Differential Adhesion

Our results can be intuitively understood in the following way: all properties of the model are fixed except the cost of a cell edge, corresponding to adhesion (Steinberg, 1962), or surface tension (Harris, 1976; Brodland and Chen, 2000). This means that the energy cost per unit area of the cell being a certain size is fixed and we are varying the cost per unit length of the cell boundary; we are adjusting the relative cost of optimising the ratio of surface area to volume.

If the cost of edges is high, there is a strong motivation to have as few as possible. If the cost of heterotypic edges is high compared to homotypic edges, there will tend to be fewer of the former and more of the latter. The pattern formed should then be the one that encloses the largest possible homotypic region: ideally a circle. This was observed by Cachat et al. (2016) when they used very unequal ratios of E-and P-cells: the less populous type formed rounded islands in a sea of the other cell type (we have also also reproduced this result in simulation by the same techniques described here: data not shown). By symmetry, the same can be expected regardless of which clone is less populous. When they are equal in number as they are in the experiments and simulations described here, however, both populations try to form circular patches. It is not possible to tile a plane with circles, so we observe elongated patches in both simulation and reality.

If the cost of heterotypic and homotypic edges are similar, no special pattern arises because there is no energy benefit to having fewer of one than the other. Therefore patterning is most likely to happen, and happen quickly, the greater the difference. In the presented model, there is a limit on how great this difference can be. The constraint 2Λ_*xx*_ *>* Λ_*xy*_ means that it must be more expensive to have two homotypic edges than on heterotypic one. Without this, the integrity of the tissue is lost as it tears itself apart wherever a heterotypic edge is incident on a boundary vertex.

### 2.5. Non-Destructive Measurement of Edge Cost

Current methods for measuring adhesivity are either destructive (e.g. laser ablation (Farhadifar et al., 2007)) or perturb the cells (e.g. tapping-mode atomic force microscopy). It seems likely that, with proper calibration, measurements of the rate of pattern formation using signal processing techniques described here, can be used to infer adhesivity or edge elasticity from time-lapse series of images of a straightforward 2-D culture of a cells mixture. This can provide developmental biologists with insight into the biophysical properties of their cells, and tissue engineers with clear information about whether self-organization of an organoid (Unbekandt and Davies, 2010; Lefevre et al., 2017) is likely to take place.

## 3. Methods

### 3.1. Cell culture and imaging

Cell images used for the comparison of phase separation patterns in real cells and in our model were acquired during a study we have described in detail elsewhere (Cachat et al., 2016). Briefly, two distinct HEK cell populations, engineered with synthetic constructs driving the expression of E-or P-cadherin adhesion molecules, were mixed with a 50/50 ratio after E-cells were marked with a green fluorescent dye. 24 h after inducing cadherin expression with tetracycline, cell monolayers were fixed, DAPI stained and imaged with a confocal microscope. Images were analysed on imageJ where DAPI nuclei were detected and assigned an identity (E-or P-nuclei) by measuring green fluorescence intensity at corresponding coordinates.

### 3.2. Mechanical model

The basis for our model of tissue dynamics are the established methods methods (Nagai and Honda, 2001; Farhadifar et al., 2007) featuring a graph (vertices, edges) describing the layout of the tissue, topological transitions (see Rules 1) for changing the layout and a potential energy function, Equation 2, from which the forces present in the system can be derived using a directional gradient, *∂* _*i*_. This is the Farhadifar potential,

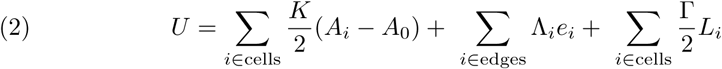

where *A*_*i*_ is the area of the *i*th cell and *L*_*i*_ is its perimeter. The *e*_*i*_ are the length of each edge and in our implementation the Λ_*i*_ depend on whether the edge is heterotypic or homotypic and of which type. See the Supplementary Materials, Section A.1 for a more detailed account.

### 3.3. Noise

Following Osborne et al. (2017) we add noise to the standard formulation of the vertex model. The addition of noise to the forces applied to the vertices is implemented discretely for each time-step *δt*, the position of each vertex being changed according to a Wiener process or Brownian motion of magnitude *D*.

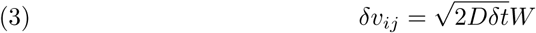

This additive noise is necessary so that the system does not become stuck in a locally stable but likely uninterestingly patterned configuration and instead moves towards a global optimum.

### 3.4. Computer simulation

We have implemented the above model using the Chaste platform (Mirams et al., 2013). This differs from the so-called “quasistatic” method of Farhadifar et al. (2007) in that instead of directly minimising the potential in Equation 2, the simulation progresses by numerically integrating the (differential) equations of motion derived from the potential. This approach affords the possibility of introducing the necessary noise described above, or indeed other arbitrary forces. Since the potential-derived force sets the tissue in motion, in order to arrive at a static equilibrium, a velocity-dependent dissipative force is needed, and indeed this is furnished for us by the Chaste environment.

Our implementation^5^, together with changes needed to the Chaste framework itself ^6^ are available on GitHub.

### 3.5. Measures for patterning

Having produced a prodigious quantity of simulation data for various adhesivity values, we needed to quantify the rate of pattern development and the degree of similarity with the patterns from the wet-lab experiments. In order to be as generally applicable as possible, to be able to calculate these measures even when cell adjacency data is noisy or unavailable, we turned to a pair of techniques from machine learning and signal processing. We used two unrelated methods in order to increase the robstness of our results. While general and not requiring anything more than images, both are sensitive to scale and resolution and require that the images to be compared are similarly calibrated with the same number of pixels per cell on average.

#### 3.5.1. Neighbour Entropy

The first measure is entropy-based. The use of entropy measures for image similarity is well established with a long history. All such measures fundamentally require a representation of the image as a probability distribution. The entropy of such a distribution is well-defined (Shannon, 2001) and from it several measures that have proven valuable in image analysis are possible such as mutual information (Smith, 1964) and the Kullback-Leibler divergence (Kullback and Leibler, 1951). We define the probabilty distribution on an image in A.6 in such a way as to capture the clustering typical of the observed patterns. It turns out that it is possible to glean the relationship between patterning and adhesivity directly from the entropy, *H*_*N*_, without requiring recourse to the similarity (or dissimilarity) measures.

#### 3.5.2. Spectral Distance

The second measure is constructed around the observation that the notion of pattern and periodicity are closely related. Indeed the nervous system seems to contain structures optimised for recognising periodoc patterns (Campbell and Robson, 1968). Given that this is the case it would not be surprising if the representation of an image in terms of its spatial frequencies would be a succinct way of describing the pattern. The spectrum is clearly not a simple scalar value that can be said to be characteristic, so we take the approach of measuring the distance to an “obviously” patterned reference image (using a random image is equally possible, see Supplementary Text, Section B.2).

Reasoning that nothing of the mathematical model is orientation-dependent, the pattern that is produced ought to be isotropic so averaging the spectrum azimuthally about the origin is justified. We use the We use the Kantorovich-Rubinstein-Wasserstein metric (Kantorovich, 2006), *W*_2_ for this purpose as implemented in the OpenCV library (Bradski, 2000), first normalising the spectra so that they have unit integral. We denote by *D*_*S*_ the distance from a given image to our exemplar.

## Acknowledgements

W.W., M.C. acknowledge the support from the Engineering and Physical Sciences Research Council (EPSRC) grant EP/J02175X/1 and from UK Research Councils Synthetic Biology for Growth programme. W.W. also acknowledges support from the National Academies Keck Futures Initiative of the National Academy of Sciences award number NAKFI CB12. J.A.D. and E.C. acknowledge the support from the Biotechnology and Biological Sciences Research Council (BBSRC) grant BB/M018040/1 and the Leverhulme Trust grant RPG-2012-558. The authors wish to thank N. Behr, R. Farhadifar, A. Fletcher and G. Plotkin for helpful discussions.

## Data availability

The simulation data and scripts to process it in order to arrive at the results elaborated herein are available at tbd Edinburgh Datashare.

## Appendix A. Supplementary Text

### A.1. On the Vertex Model

While topological considerations tell us about the structure of an epithelial sheet, and how the structure *can* change, more information is needed to know how it *does* change. In addition to the graph of vertices and edges, to know the actual configuration of a tissue in the plane, we need the positions of these vertices. To know how the tissue changes from one moment to the next, we must calculate the forces on each vertex.

The mechanical properties of cells are precisely what distinguishes them from soap bubbles, in particular while soap bubbles are filled with a compressible gas, biological cells are, to a good approximation filled with an incompressible fluid. In both cases the surface of the cell or soap bubble has a certain tension that causes, all else being equal, the edges to be as short as possible. These observations led to the definition of a potential that captures these competing effects of internal pressure and surface tension (Nagai and Honda, 2001), later refined (Farhadifar et al., 2007) to include a second order correction that considers the role of cell perimeter in addition to individual edges.

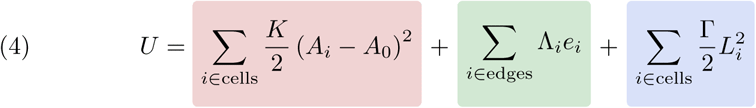

In this paper, we use the Farhadifar potential, Equation 4, normalised so that cells have a unit target area. Each edge has an associated cost, Λ_*i*_ *∊* {Λ_*P P*_, Λ_*P E*_, Λ_*EE*_, Λ_*EP*_} accordingly as it is heterotypic or homotypic for each kind of cell. Calculating the force on a vertex due to this potential is then a matter of taking the gradient of this potential along each coordinate of each vertex,

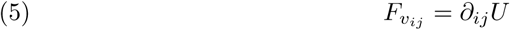

where *v*_*ij*_ is the *j*-th coordinate of the *i*-th vertex and*∂* _*ij*_ is the derivative along that axis.

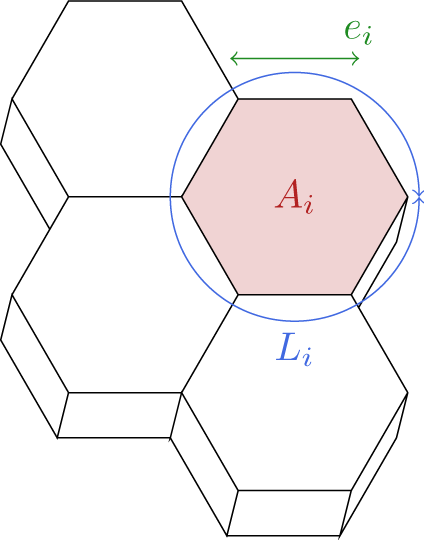

### A.2. Quasi-Static vs. Continuous Simulation

We know from the Lagrange formulation of classical mechanics how to derive, in general, the equations of motion for a system given a description of potential and kinetic energies in terms of generalised coordinates, *q*_*i*_ Goldstein (1980),

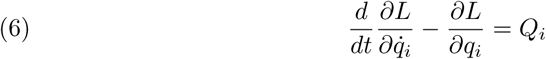

where *L* = *T - U* is the Lagrangian, made from the kinetic energy, *T* and the potential energy *U*, and the *Q*_*i*_ are *external* forces not captured by the energy description. In our case the generalised coordinates, *q*_*i*_, correspond to the coordinates of the vertices of the tissue.

For the *quasi-static* technique, as in Farhadifar et al. (2007), we suppose that there are no external forces and the system moves quickly enough to a static equilibrium configuration that transient states can be neglected. Static equilibrium means that there is no change in configuration with time, hence,

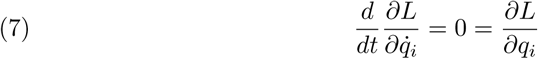

Furthermore, since in static equilibrium the system is motionless, we have *T* = 0, and all that is necessary is to find *q*_*i*_ such that,

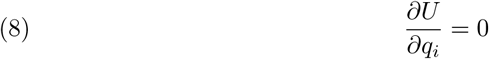

This approach has a disadvantage in that an equilibrium, or local minimum of *U* will be found (if it exists), it is unspecified *which* minimum it is, and in particular if it is the same one which would be reached from a previous non-equilibrium state following a perturbation such as mitosis. If not done carefully, the configuration of the tissue may jump between plausible though unrelated equilibrium configurations which though they are sufficient to recover statistics about polygon distribution do not necessarily represent a realistic development of the tissue.

For this reason in the present paper we have adopted the approach using the Chaste software (Mirams et al., 2013) which integrates Equation 6 directly (see (Fletcher et al., 2013) for more details). This requires an external velocity-dependent force, friction or drag for the following reason. Suppose *Q*_*i*_ = 0, no such external forces. Consider a system that is out of equilibrium where 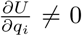. With a little algebra, or simply by noting that the total energy, *T* + *U*, must be conserved, some infinitesimal time later, the kinetic energy *T* must be non-zero. Eventually a state is reached where 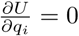 but now the other term,

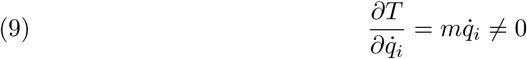

since the system is in motion. This momentum carries it past what would otherwise have been a potential minimising equilibrium configuration. The role of the external force is to remove this excess energy from the system and allow it to settle to a stable, static equilibrium configuration.

### A.3. Image Segmentation

A threshold function is used to assign each pixel a black or a white value depending on which cell type it is:

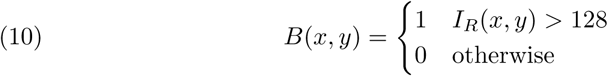

where *I*_*R*_(*x, y*) is the red pixel value for position (*x, y*) in the image and can take on values in the range [0, 255], as in,

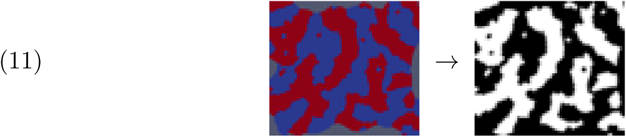

### A.4. Lattice Reconstruction

The confocal imagery from experiments *in vitro* required some special handling in order to be able to apply the coarse segmentation technique of Section A.3. This was due to the fact that the processing using the ImageJ software produced cell centroids but not the cell boundaries necessary for colouring in to make a binary image of the pattern. To solve this we used the technique of Jones et al. (2005) of constructing a Voronoi tesselation using these centroids. Accordingly as each centroid or cell nucleus was said to be above or below a certain thresold of green fluorescence, the corresponding Voronoi cell was coloured red or blue. Processing then proceeded as for the images from the simulation data.

### A.5. Spectral Distance

The spatial frequency spectrum distance method is based on the observation that there appears to be a periodicity in the image where a pattern is present, and uses a Fourier transform. Let *S*(*θ, r*) be two dimensional fourier transform of the input image *B*(*x, y*) parametrised by angle and distance from the origin (Figure 2, coloured panels). We define a one-dimensional azimuthally averaged spectrum (Figure 2, graphs),

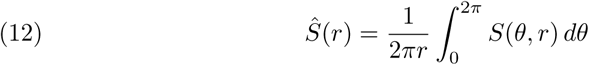

From Equation 12 we define a spectrum distance metric for comparing images to a given reference image,

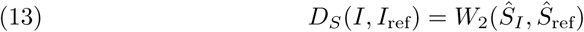

in other words the spectrum distance, *D*_*S*_ between an image and the reference (i.e. patterned) image is the Kantorovich-Rubinstein-Wasserstein distance, also known colloquially as the Earth Mover’s distance, between their azimuthally averaged spectra.

For the purposes of this paper, we use as our standard reference image, Fig 5, the clear and stable pattern produced after 1400 time units of the simulation with, Λ_*EE*_ = Λ_*P P*_ = 0.05 and Λ_*P E*_ = Λ_*EP*_ = 0.096, or in other words, near optimal parameter values for obtaining patterning. For brevity, where there is no risk of ambiguity, we write,

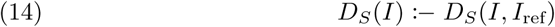

for this reference image.

**Figure 5.**
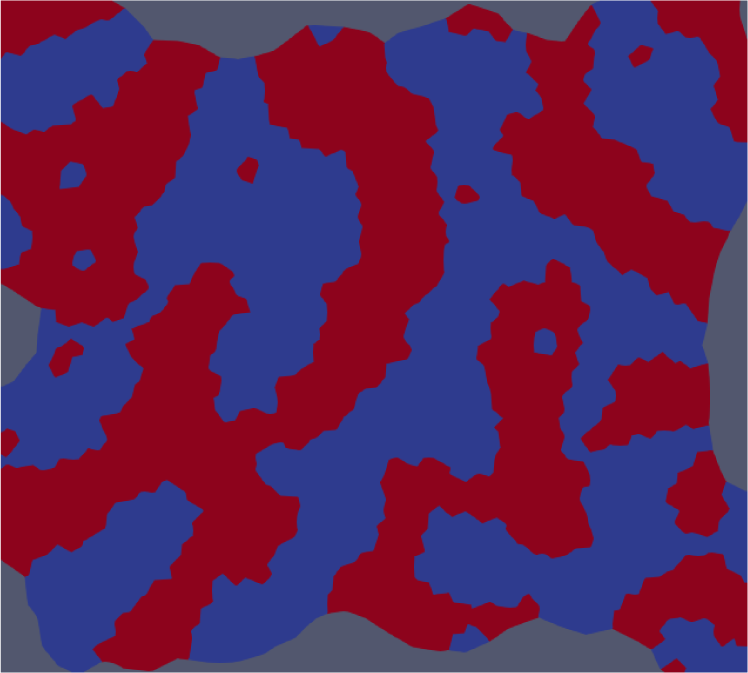
Standard patterned reference image

## A.6. Neighbour Entropy

The neighbour entropy metric recognises that the basic process at work in patterning is minimising the total length of heterotypic edges. Given a lattice of pixels or cells (this measure is meaningful in either case), we first define the pair of probabilities, *P*_*E*_ and *P*_*P*_, for a randomly selected cell being is *E* or *P* type. Further, we define the conditional probabilities, *P*_*EE*_, *P*_*EP*_, *P*_*P E*_, *P*_*P P*_ that, supposing a cell is of *E* type, a randomly chosen neighbour is of type *E* or *P*, *mutatis mutandis* for the other cases. Combining these, we obtain the following distribution describing a population of cells, and their neighbours:

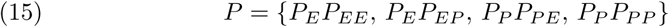

This distribution gives rise to an entropy measure,

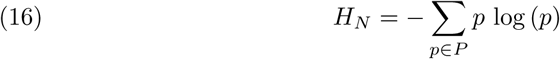

which can be calculated for any image.

## Appendix B. Supplementary Data

### B.1. Fixed homotypic edge cost

In the main text we fixed the heterotypic edge cost, Λ_*P E*_ = Λ_*EP*_ = 0.096 and allowed the heterotypic edge cost to vary. Here, for completeness, we show the reverse in Fig 6. As would be expected, the pattern develops most quickly (the spectral distance to the patterned reference image decreases most rapidly, as does the neighbour entropy) for those values that maximise the difference between the heterotypic and homotypic edge costs.

**Figure 6.**
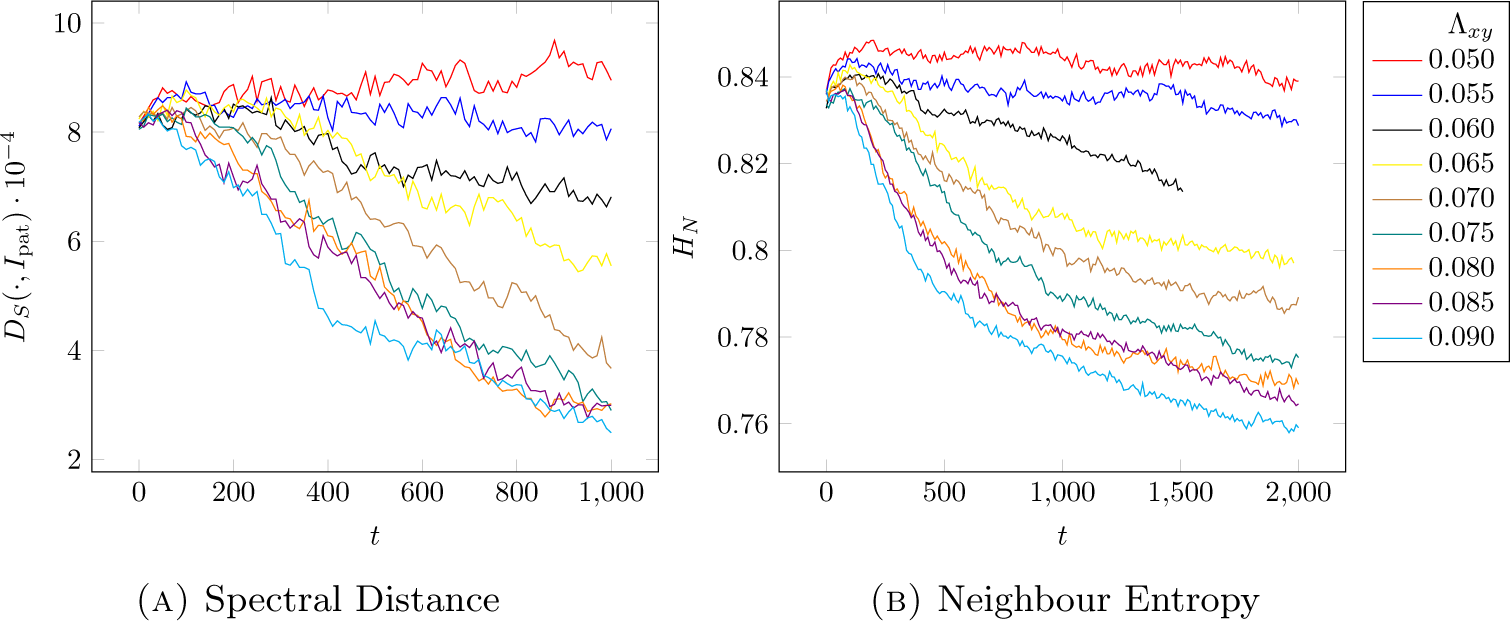
Spectral distance and neighbour entropy for traces for various values of Λ_*P E*_ = Λ_*EP*_. In all cases the heterotypic value is Λ_*P P*_ = Λ_*EE*_ = 0.05.

### B.2. Spectral distance to random configuration

For the most part we have chosen to measure the spectral distance to an exemplar highly patterned image. This is partly because it was already established in Cachat et al. (2016) that the observed patterning is non-random and the goal here is to establish a technique for navigating a richer landscape of patterns endowed with a metric. It is perfectly reasonable instead to ask what the distance from the random initial configuration is; to use *D*_*S*_ (*I, I*_initial_) rather than *D*_*S*_ (*I*_ref_). Fig 7 shows what this looks like.

**Figure 7.**
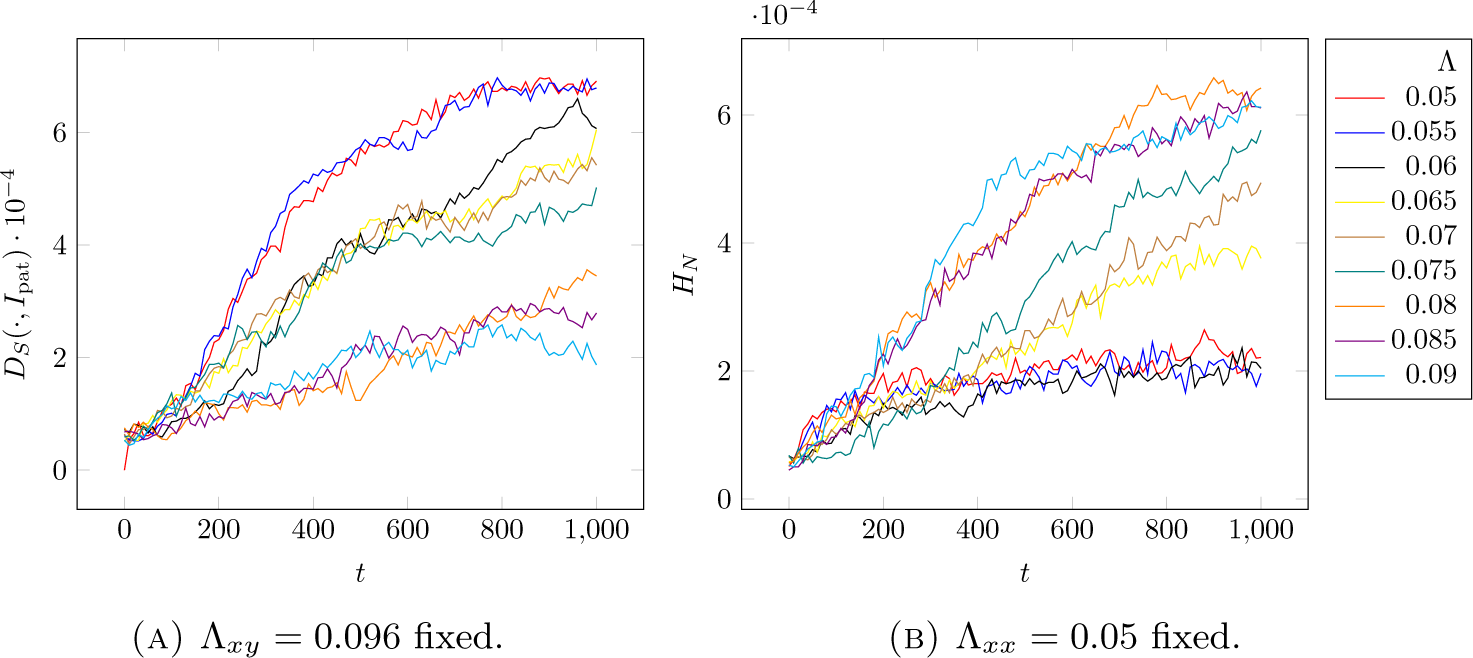
Spectral distance from initial random configuration, holding either heterotypic or homotypic edge cost fixed.

In time, the system moves away from the initial conditions. The distance increases dramatically more quickly for values of Λ where patterning is observed. Even for values of that do not cause patterning the distance increases, due to noise and the fact that the initial conditions are artificially regular. This tells us, interestingly, that we are exploring distance in a much more compex space than a simple one-dimensional, “pattern or no pattern". It is possible to move away from a configuration of “no pattern”, to see distance increase, and yet not move closer to a patterned configuration. This in turn suggests that it was the correct choice to use distance from a patterned image to illustrating the phenomenon of pattern development that we wished to show.

### B.3. Integrity constraint violation

The integrity constraint that says that it must be more expensive to have two edges at the boundary of the tissue than to have one heterotypic edge,

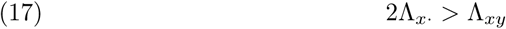

combined with our convention to use the *same* value for these edges as for homotypic edges, Λ_*x·*_ = Λ_*xx*_, is required to avoid the situation shown in Fig 8. The subscript *x* is intended to take on either of the values *P* or *E*. If the constraint is violated, any heterotypic edge incident on the tissue boundary is liable to split into two, cheaper, edges, and the tissue comes apart at the seams, with homogeneous islands detatching themselves from the main tissue.

**Figure 8.**
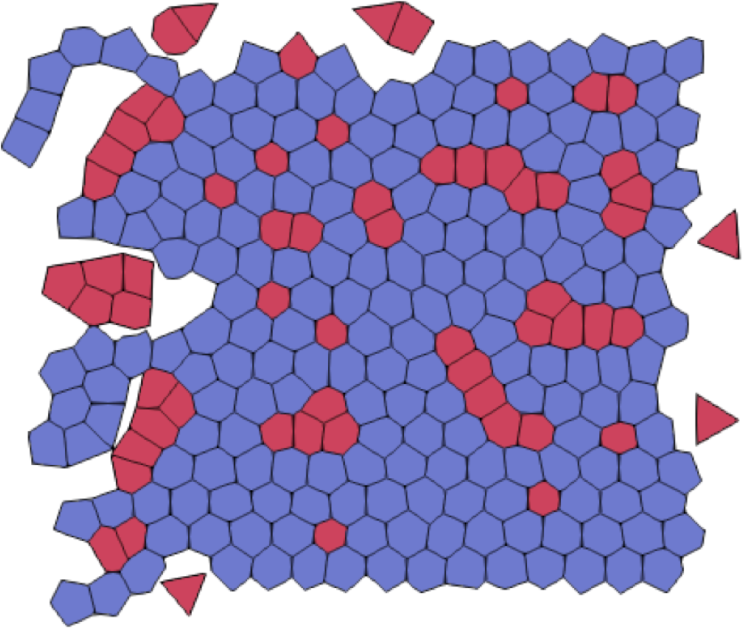
Integrity constraint violation

### Footnotes

5 https://github.com/edinburgh-rbm/chaste_tissue

6 https://github.com/wwaites/chaste

